# Dissecting the collateral damage of antibiotics on gut microbes

**DOI:** 10.1101/2020.01.09.893560

**Authors:** Lisa Maier, Camille V. Goemans, Mihaela Pruteanu, Jakob Wirbel, Michael Kuhn, Elisabetta Cacace, Tisya Banerjee, Exene Erin Anderson, Alessio Milanese, Ulrike Löber, Sofia K. Forslund, Kiran Raosaheb Patil, Georg Zeller, Peer Bork, Athanasios Typas

## Abstract

Antibiotics are used for fighting pathogens, but also target our commensal bacteria as a side effect, disturbing the gut microbiota composition and causing dysbiosis and disease^1-3^. Despite this well-known collateral damage, the activity spectrum of the different antibiotic classes on gut bacteria remains poorly characterized. Having monitored the activities of >1,000 marketed drugs on 38 representative species of the healthy human gut microbiome^4^, we here characterize further the 144 antibiotics therein, representing all major classes. We determined >800 Minimal Inhibitory Concentrations (MICs) and extended the antibiotic profiling to 10 additional species to validate these results and link to available data on antibiotic breakpoints for gut microbes. Antibiotic classes exhibited distinct inhibition spectra, including generation-dependent effects by quinolones and phylogeny-independence by β-lactams. Macrolides and tetracyclines, two prototypic classes of bacteriostatic protein synthesis inhibitors, inhibited almost all commensals tested. We established that both kill different subsets of prevalent commensal bacteria, and cause cell lysis in specific cases. This species-specific activity challenges the long-standing divide of antibiotics into bactericidal and bacteriostatic, and provides a possible explanation for the strong impact of macrolides on the gut microbiota composition in animals^5-8^ and humans^9-11^. To mitigate the collateral damage of macrolides and tetracyclines on gut commensals, we exploited the fact that drug combinations have species-specific outcomes in bacteria^12^ and sought marketed drugs, which could antagonize the activity of these antibiotics in abundant gut commensal species. By screening >1,000 drugs, we identified several such antidotes capable of protecting gut species from these antibiotics without compromising their activity against relevant pathogens. Altogether, this study broadens our understanding of antibiotic action on gut commensals, uncovers a previously unappreciated and broad bactericidal effect of prototypical bacteriostatic antibiotics on gut bacteria, and opens avenues for preventing the collateral damage caused by antibiotics on human gut commensals.

## MAIN TEXT

Medication is emerging as major contributor for changes in the composition of the human gut microbiota^4,13-15^. Such severe and long-lasting changes are associated, and in some cases causatively linked, to dysbiosis and a wide range of diseases^16^. Although several non-antibiotic drugs may also have a previously unappreciated impact on the gut microbiome composition^4,16,17^, antibiotics, developed to have broad spectra and thereby target very diverse pathogens, are long known to take a heavy toll on our gut flora, causing a variety of gastrointestinal side-effects^18^, including *Clostridioides (former Clostridium) difficile* infections. Recently more attention has been given to this collateral damage of antibiotics on the gut microbiota and thereby on the host’s wellbeing. *In vivo* studies highlight links between the long-term microbiota compositional changes and host dysbiosis, including the development of allergic, metabolic, immunological and inflammatory diseases^5-8,10,11,19-21^. While uncovering the direct effects of different antibiotics on our gut flora is critical to improve general health, technical difficulties hamper routine testing of antibiotic susceptibility in anaerobes^22,23^. Currently available data on bacterial susceptibility to antibiotics is focused on invasive pathogens and offers little to no resolution in the diversity of the human gut microbiota^24^. Information is missing even for the most prevalent and abundant gut species, or ones recently associated with dysbiosis and disease^25,26^. In addition, existing animal or cohort studies have used a handful of antibiotics or merge data from different antibiotic classes, precluding systematic and general conclusions on the matter.

We recently assessed the direct effect of ∼1200 FDA-approved drugs on the growth of 38 prevalent and abundant or disease-associated human gut species under anaerobic conditions at a fixed concentration of 20 µM^4^. This initial screen (referred to hereafter as “screen”) included 144 antibiotics (Fig. 1a, Extended Data Fig. 1, Suppl. Table 1), with different classes having discernible effects on gut microbes (Fig. 1b). We validated these results by measuring 815 MICs (33 antibiotics and 2 antifungals for 17 species, 22 antibiotics for 10 additional species), using MIC gradient test strips (Fig. 1a, Extended Data Fig. 2, Suppl. Table 2 + 3). Despite differences in the experimental procedure, concordance between data from the initial screen and MICs is very high: Specificity and sensitivity of 0.97 (Extended Data Fig. 3a). The newly established MICs also correlate well with available data on antimicrobial susceptibility from databases such as EUCAST^24^ or ChEMBL^27^ (r_s_=0.69 and r_s_=0.64, respectively), despite differences in strains and media used (Extended Data Fig. 3b). Importantly, this new dataset considerably expands the available MICs, as much as by 80% for non-pathogenic bacteria (Fig. 1c, Extended Data Fig. 3c). Altogether, the initial screen and the new MIC dataset provide high-resolution information on the target spectrum of antibiotics on commensal gut microbes, which we explored further.

**Figure 1.**
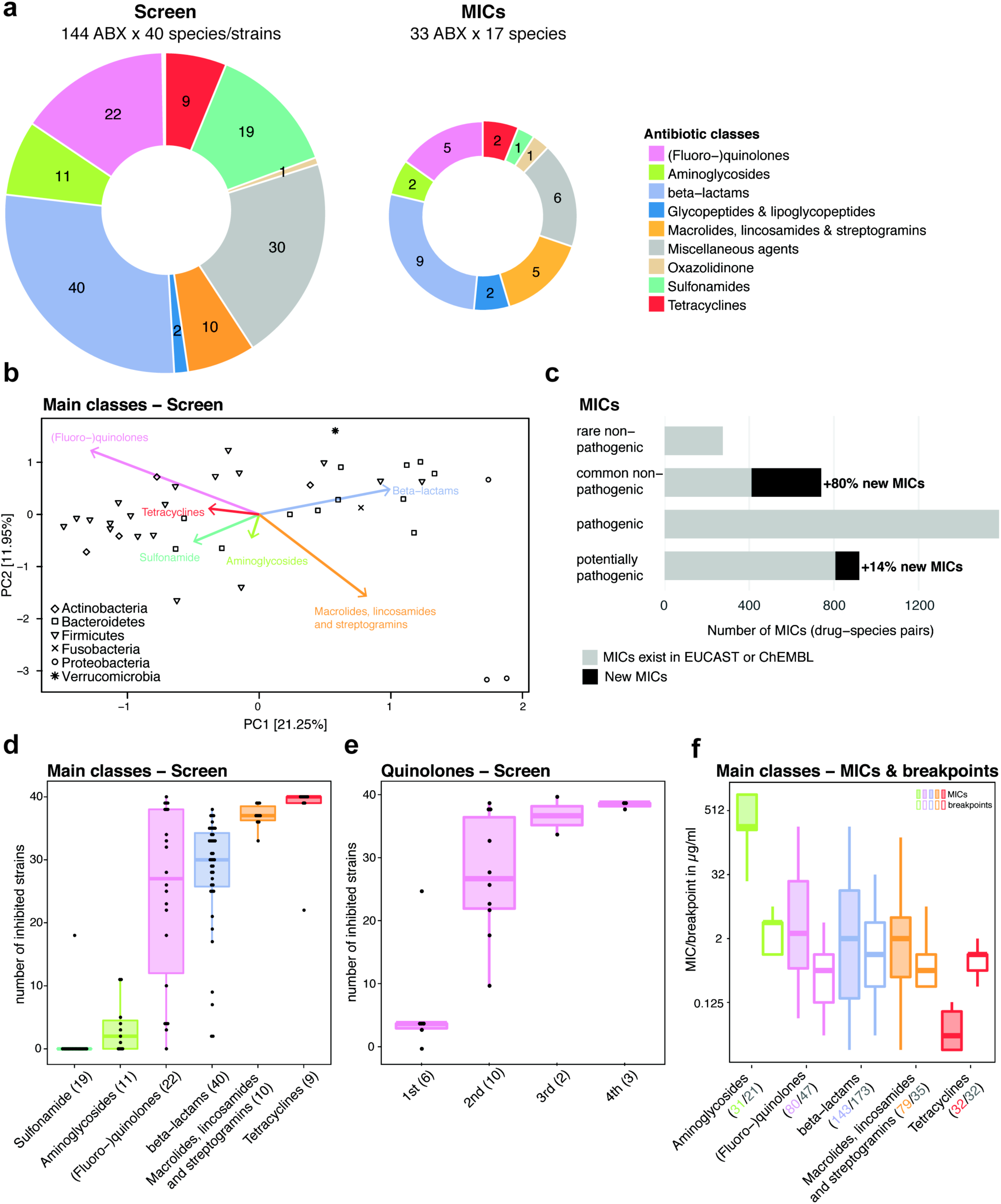
Activity spectrum of antibiotic classes on human gut commensals. **a**. Overview of antibiotics tested in initial screen at 20 µM concentration^4^ and validated by MIC determination in this study. **b**. Principal component analysis based on AUCs from the initial screen on the effects of antibiotics on gut commensals. Antibiotic classes drive some separation at the phylum level, e.g. beta-lactams separate Bacteroidetes and macrolides/lincosamides/streptogramins separate Proteobacteria. **c**. Comparison of MICs from this study to MICs available from public databases. Species are classified as “common” or “rare” if they are present in the gut microbiome of more or less than 1% of 727 healthy individuals, respectively (see Methods). **d**. For the main antibiotic classes from the screen, the numbers of inhibited strains are shown (N as in **a**). 40 strains tested in total at a 20 µM antibiotic concentration. Boxes span the IQR and whiskers extend to the most extreme data points up to a max of 1.5 times the IQR. **e**. Number of inhibited strains per (fluoro-)quinolone drug generation. Number of tested drugs per generation is indicated in brackets on x-axis labeling. Boxplots as in panel **d. f**. MICs of drug-species pairs for the main antibiotic classes measured in this study are depicted next to EUCAST clinical (susceptibility) breakpoints for pathogens. Numbers of drug-species pairs (MICs; colored) and of antibiotic per class (EUCAST clinical breakpoints; grey) are shown in brackets. Boxplots as in panel, **d**, y-axis is log2 scale.

The antibiotics tested exhibited strong class-dependent effects (Fig. 1b, d). Consistent with literature, aminoglycosides hardly affected gut microbes under anaerobic conditions^28^ and sulfonamides were inactive in the medium used for the screen^4^. Quinolones acted in a generation-dependent manner. First-generation variants were effective only on a narrow spectrum of microbes that included both commensal *E. coli* tested. Second- and third-generation quinolones increased the spectrum. Fourth-generation variants (developed to increase activity against anaerobes) inhibited all tested species, except for *Akkermansia muciniphila* (Fig. 1e, Extended Data Fig. 1, red box), a species associated with protection against different diseases and dysbiotic states^29^, and even positive responses to immunotherapy^30^. For β-lactams, resistance was patchy but distinct for different members and subclasses (Extended Data Fig. 2, 4a). For *Bacteroidetes*, we tested additional species and strains (in total 12 and 19, respectively) (Extended Data Fig. 4b, c), confirming that β-lactam sensitivity and phylogenetic relatedness are uncoupled (Extended Data Fig. 4d). This argues for resistance mechanisms being strain-specific and horizontally transferred. Macrolides showed a strong impact on gut commensals and inhibited all tested microbes (Fig. 1d), except for the opportunistic pathogen *C. difficile*, which was resistant to all tested macrolides and clindamycin (Extended data Fig. 2, red box). This is in line with the associated risk of *C. difficile* infection after macrolide/clindamycin treatment^31^. Finally, 8 of the 9 tested tetracyclines inhibited nearly all tested microbes, which is surprising in the light of the gut microbiota being considered as reservoir for tetracycline resistance genes^32^.

Concentration-resolved MICs confirmed the same drug class-dependent trends observed in the screen (Fig. 1d, f). In addition, MICs allow for comparisons with clinical breakpoints, i.e. MICs at which a species should be considered resistant or susceptible (Fig. 1f). Overall, the gut microbes in our assays (anaerobic growth, gut mimetic growth medium^33^) were slightly more resistant to most antibiotic classes than previously reported for pathogens (aerobic growth, Mueller-Hinton agar). Tetracyclines were the exception, inhibiting commensals at significantly lower concentrations than what is reported for pathogens (Fig. 1f). Thus, commensals might be considerably less resistant to tetracyclines than previously anticipated and suggested by the detection of tetracycline resistance elements in fecal metagenomes.

Recent *in-vivo* studies have shown that β-lactams and macrolides have a strong and long-lasting collateral impact on the gut microbiota composition and thereby on host health^5-8^. As β-lactams exhibited strain-specific effects (Extended Data Fig. 1, 2, 4) and are known to kill bacteria (bactericidal), they could *irrevocably* deplete *specific* members of the gut microbiota, thereby explaining their differential and long-lasting effects on the community composition. On the other hand, macrolides uniformly targeted all tested gut commensals (Fig. 1d) and are textbook bacteriostatic antibiotics, i.e. inhibit bacterial growth, but do not kill (at least at high numbers). In this case, the long-term community composition change is more difficult to rationalize, as all community members are inhibited, but should be able to regrow once drug is removed. Similarly, tetracyclines, another class of bacteriostatic antibiotics that acted on nearly all gut microbes we tested, have known gastro-intestinal side-effects^18^, which are indicative of gut microbiome dysbiosis. We thus wondered at which level macrolides and tetracyclines exert a differential effect on gut microbes. Although traditionally both clinical use^34-37^ and basic research^38,39^ heavily rely on this distinction between bactericidal and bacteriostatic antibiotics, there are reports of drugs changing their killing capacity depending on the organism, drug concentration or medium tested^40,41^ (and increased evidence from meta-analyses that the distinction may have little relevance to clinical practice^42,43^). We therefore hypothesized that this bacteriostatic/bactericidal divide may be less rigid for gut commensals, which are more phylogenetically diverse than the few pathogens usually tested, and hence provide a level where the effect of these drug classes on gut microbes becomes differential.

The standard way to determine whether an antibiotic has bactericidal or bacteriostatic activity is to calculate time-kill curves, where the bacterial survivors are counted on agar at various time-points after drug treatment. If, over a significant period of antibiotic treatment (ranging from 5 to 24 hours), the number of colony forming units (CFU)/ml of culture decreases by more than 99.9%, the antibiotic is considered bactericidal^40^. We assessed the survival of 12 abundant gut microbes over a 5-hour treatment of either a macrolide (erythromycin or azithromycin) or a tetracycline (doxycycline) at 5 x MIC (Fig 2a + b, Extended Data Fig. 5). About half of the tested species decreased in survival by >99.9%, pointing to these drugs being bactericidal to several abundant gut microbes. To confirm this further, we tested the viability of *B. vulgatus* and *E. coli* ED1a upon erythromycin, azithromycin or doxycycline treatments using live/dead staining. Microscopy and flow cytometry assessment of live/dead bacteria corroborated the initial observations (Fig. 2c, Extended Data Fig. 6). As tetracyclines are considered bona-fide bacteriostatic drugs in *E. coli*, we were surprised to see that doxycycline effectively killed the commensal *E. coli* ED1a (Fig. 2a). We verified that these effects held also in the presence of oxygen (Extended Data Fig. 7a) and confirmed that doxycycline has a stronger bactericidal action on this natural isolate than on the domesticated *E. coli* K-12 lab strain, BW25113 (Extended Data Fig. 7b). In parallel, we excluded that the differences in killing capacity were confounded by growth rate, growth phase or MIC of the bacterial species tested (Extended Data Fig. 8). We also noticed that *B. vulgatus* and *B. uniformis* cultures decreased density in the presence of erythromycin (Fig. 2d). We confirmed by time-lapse microscopy that this was due to lysis. Erythromycin caused cell shape defects, including blebbing, cytoplasmic shrinkage, and ultimately cell lysis in both *B. vulgatus* and *B. uniformis* (Figure 2e, Movies 1-4). Altogether, this selective bactericidal activity of macrolides and tetracyclines on specific gut commensals could provide an explanation for the strong effects these drug classes have on the gut microbiota composition of human individuals. The gut microbes killed from the drug would be inadvertently removed from community, whereas the ones being only inhibited could recover when the therapy stops.

**Figure 2.**
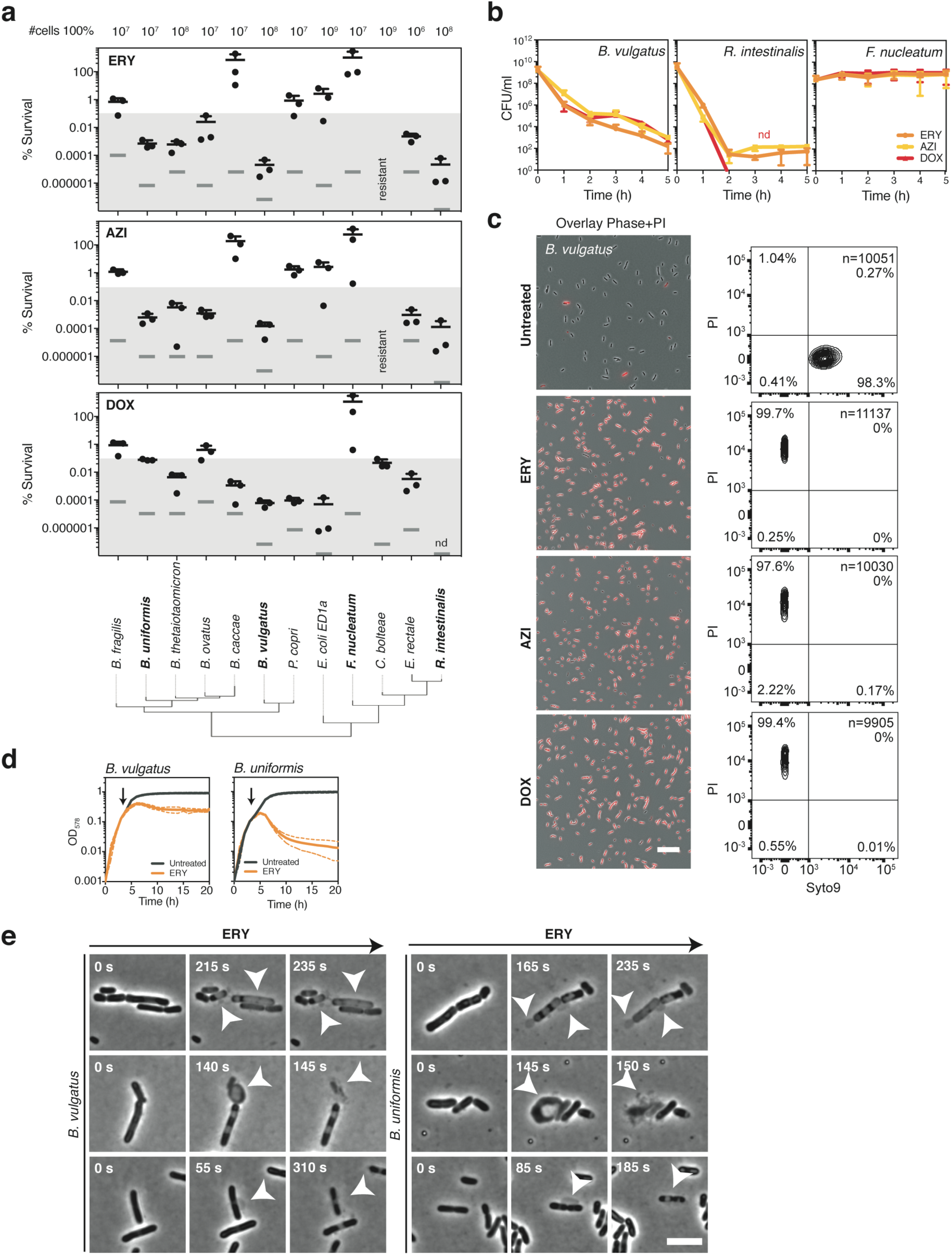
Macrolides and tetracyclines kill human gut commensal species. **a**. The survival of 12 abundant gut microbe species was measured after a 5-hour treatment with a 5-fold MIC of erythromycin (ERY), azithromycin (AZI) or doxycycline (DOX). The survival was assessed by counting CFUs/ml before and after antibiotic treatment. The number of CFUs/ml before treatment was set as 100%. The detection limit for each experiment (gray bar) and the bactericidal threshold (shaded area) are indicated. Species are plotted according to phylogeny (IQTree, Methods) and in bold are noted the species that are used in later panels. The graph shows the mean+SD of 3 independent experiments. **b**. Time-kill curves of *B. vulgatus, R. intestinalis* and *F. nucleatum* after antibiotic treatments. Survival was assessed by CFU counting over a 5 hour-treatment of ERY, AZI or DOX. This graph shows the mean±SD of 3 independent experiments. Nd: non-detectable. Time-kill curves for the other tested gut microbes can be found in Extended Data Fig. 5. **c**. Live/dead staining of macrolide or tetracycline-treated *B. vulgatus*. The left panel shows an overlay of phase contrast and fluorescence microscopy images of propidium iodide (PI)-stained *B. vulgatus* before and 5 hours after ERY, AZI or DOX treatment. Cultures were concentrated before imaging; the scale bar is 10 µm. The right panel shows the corresponding quantification of live/dead-stained cells by flow cytometry with Syto9 on the x-axis (live cells) and PI on the y-axis (dead cells). Both the total number of measured events (n) and the percentage of cells found in each quadrant are indicated. **d**. Erythromycin induces lysis of *B. vulgatus* and *B. uniformis. B. vulgatus* and *B. uniformis* were grown for 3 hours before addition (yellow) or not (black) of 15 µg/ml ERY treatment (5-fold MIC; yellow) as indicated by the arrow. Growth curves were acquired for 20 hours. This graph shows the mean±SD (dotted line) of 3 independent experiments. **e**. Erythromycin induces blebbing, cytoplasmic shrinkage and lysis in *B. vulgatus* and *B. uniformis*. Phase contrast movies of *B. vulgatus* and *B. uniformis* were acquired after ERY treatment (5-fold MIC). Here shown 3 frames of 3 images per strain (time indicated in the upper left corner; t=0 when drug added). White arrows indicate blebs, cytoplasmic shrinkage and bacterial lysis; the scale bar is 5 µm. Movies are available in Supplementary Material (Movies 1-4).

Knowing that drug combinations often have species-specific outcomes^12^, we reasoned that we could identify drugs that selectively antagonize the effect of antibiotics on gut microbes, while retaining activity against pathogens. Therefore, we screened the Prestwick library to identify antagonizing compounds to erythromycin or doxycycline on two abundant and prevalent gut microbes, *B. vulgatus* and *B. uniformis* (Fig. 3a, Extended Data Fig. 9). Of the 19 identified hits (Fig. 3b, Suppl. Table 4), we tested the 14 candidates with the strongest activity in a concentration-dependent manner (Extended Data Fig. 10a). Nine retained antagonistic activity over a broader concentration range, which we confirmed by checkerboard assays (Fig. 3c). The antidotes that showed the strongest antagonisms were the anticoagulant drug dicumarol, and two non-steroidal anti-inflammatory drugs, tolfenamic acid and diflunisal. While dicumarol rescued *B. vulgatus* from erythromycin and diflunisal from doxycycline, tolfenamic acid was able to protect *B. vulgatus* from both drugs. In addition, these interactions were able to at least partially rescue the killing of *B. vulgatus* by erythromycin and doxycycline (Extended Data Fig. 10b). We then probed two of these drugs for their ability to protect other abundant gut commensals and confirmed that both dicumarol and tolfenamic acid were able to counteract erythromycin on several species (Fig. 3d, Extended Data Fig. 11). In contrast, both drugs did not affect the potency of erythromycin on *Staphylococcus aureus, Streptococcus pneumoniae* and *Enterococcus faecium*, pathogens against which erythromycin is active/prescribed (Fig. 3e, Extended Data Fig. 12a). For example, tolfenamic acid and dicumarol at concentration ranges of 5-40 µM could rescue the growth of five out of seven tested abundant gut commensal species at clinically relevant erythromycin concentrations (Fig. 3f, Extended Data Fig. 12b). Altogether, our data provides a proof-of-principle for identifying antidotes that specifically mask the collateral damage of antibiotics on commensals. This concept would need to be further validated in the future in animal models. Antidotes may also need to be modified to late (colon)-release or non-absorbable formulations to ensure they reach the gut and to minimize adverse effects from their primary action.

**Figure 3.**
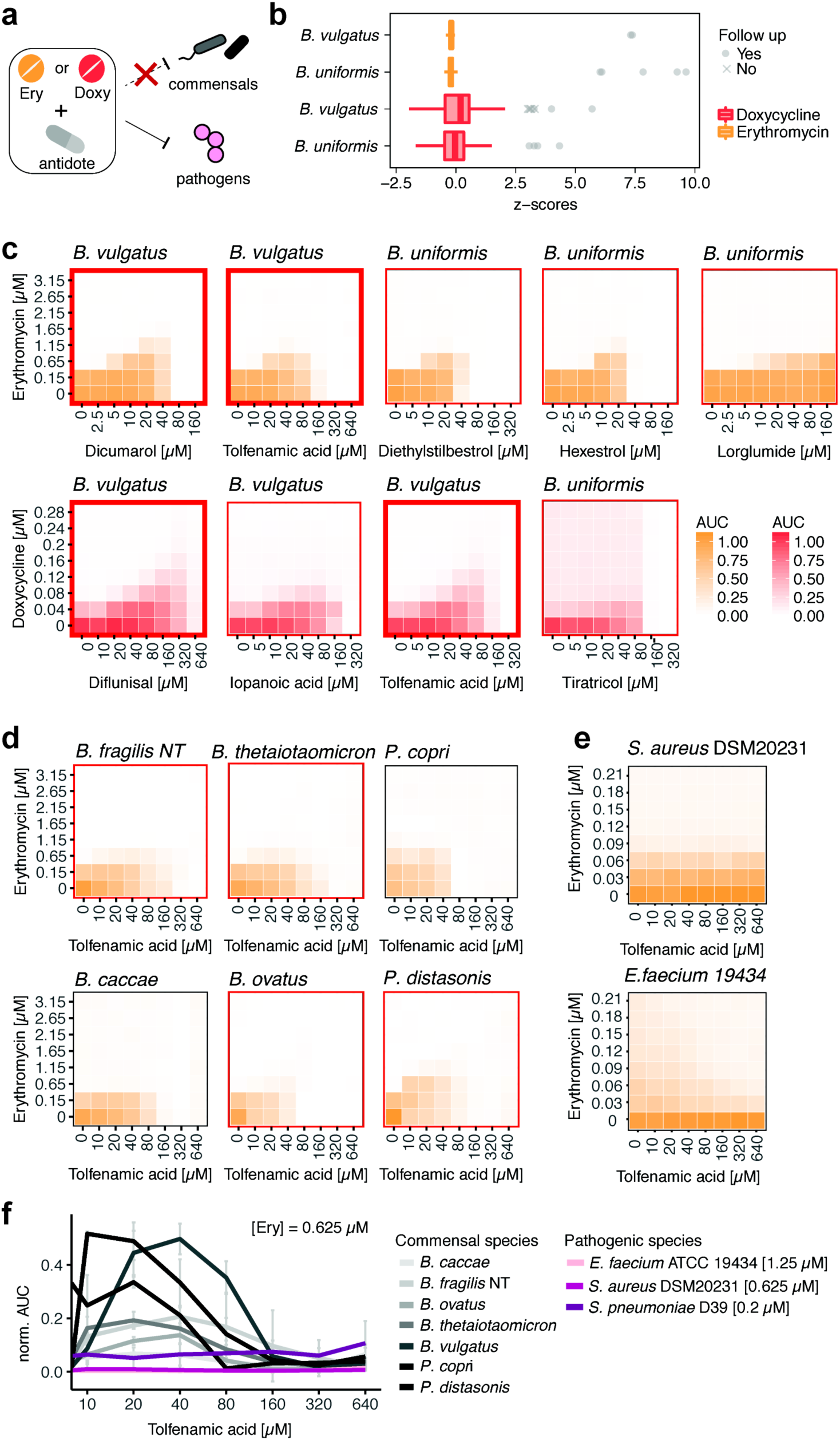
Antidotes for selective protection of prevalent and abundant gut commensal species from macrolides and tetracyclines. **a**. Schematic illustration of the screen concept: searching for antidote compounds that antagonize the antibacterial effect of erythromycin or doxycycline on commensal but not on pathogenic bacteria. **b**. Z-scores on bacterial growth (based on areas under the curve (AUCs)) for combinatorial drug exposure with antibiotic (ERY or DOX) and FDA-approved drug. Compounds that successfully rescued *B. vulgatus* and/or *B. uniformis* growth in the presence of the antibiotic (z-score > 3) are indicated in gray. The strongest hits (circles) were validated further in concentration-dependent assays (Extended Data Fig. 10a). For each antibiotic and each strain, ∼1200 drugs were tested in two replicates. Boxplots are defined as in Figure 1d. **c**. For 9 of the validated antagonists, 8 x 8 checkerboard assays were performed to determine concentration ranges of the antagonistic interaction. Heat maps depict bacterial growth based on normalized median of AUCs of 4 replicates. All interactions were antagonistic, and pairs tested further in other commensal species are framed in bold. **d**. Checkerboard assays confirm the ability of tolfenamic acid to protect further gut commensals from growth inhibition by erythromycin. Heat map as in **c**, but for 2 replicates. Antagonistic interactions are framed in red. **e**. Checkerboard of tolfenamic acid with erythromycin reveal neutral interactions in *S. aureus* and *E. faecium* (aerobic conditions). Heat maps as in c, based on at least two independent experiments with two technical replicates each. **f**. Tolfenamic acid concentration-dependent rescue of commensal growth at clinical relevant erythromycin concentrations based on AUCs (anaerobic conditions). Erythromycin still retains its activity against pertinent pathogens such as *S. aureus, E. faecium* and *S. pneumoniae* (aerobic conditions). 0.625 µM correspond to ∼0.5 µg/ml erythromycin, which is in the range of the MIC breakpoints for *Staphylococcus* (1 µg/ml), *S. pneumoniae* (0.25 µg/ml) and *Streptococci* groups A, B, C & G (0.25 µg/ml). Error bars depict standard deviation.

In summary, our study provides a high-resolution map of the collateral damage of antibiotics on 50 different resident gut microbes down to the level of individual drugs, species and partially even strains. We challenge the universal divide of antibiotics into bacteriostatic and bactericidal across bacteria, as this breaks down when tested beyond model organisms. Antibiotics with preferential killing of some species may be the most detrimental to our gut flora, although the first studies in a few healthy individuals point to the gut microbiota having some resilience against specific antibiotic regimens^44^. Understanding the underlying mechanisms for this selective killing might open up ways for the development of new antimicrobials, but also strategies for controlled microbiome modulation^15^. Finally, we provide a proof-of-concept that species-specificity of drug combinations^12^ can be exploited to identify antidotes that selectively protect the gut microbiota from the adverse effects of systemic antibiotic therapy. This new approach adds to proposed and existing strategies of gut microbiota protection against antibiotics, such as co-administration of activated charcoal^45^, β-lactamases^46^, probiotics or (autologous) fecal transplants^47^. Overall, our results suggest that interactions of antibiotics and commensals merit deeper exploration, as our current knowledge of the mode(s) of action of antibiotics in model pathogens is not necessarily transferable to commensals.

## METHODS

### Growth conditions

All experiments from this study were performed in an anaerobic chamber (Coy Laboratory Products Inc) (2% H_2_, 12% CO_2_, 86% N_2_) and all materials and solutions used for these experiments were pre-reduced for at least 24 h before use unless specified otherwise. Bacteria used in this study were typically pre-cultured for two overnights: Cells were cultured in 5 ml modified Gifu Anaerobic Medium broth (MGAM) (HyServe GmbH & Co.KG, Germany, produced by Nissui Pharmaceuticals) and grown at 37°C overnight. The next day, cells were diluted 1/100 in 5 ml MGAM medium and grown at 37°C for a second overnight before starting the experiments.

### Quantitative assay for minimum inhibitory concentration determination with MICs test strips

MICs test strips were purchased from Liofilchem or Oxoid (Suppl. Table 2). All MICs were measured under anaerobic growth conditions inside a Coy anaerobic chamber. Bacteria were precultured in MGAM for two overnights and cultures were diluted to OD_578_ = 0.5. 50 µl of the diluted culture were spread on a MGAM agar plate and allowed to dry for 15 min. The MIC test strip was placed on the agar with sterile tweezers, allowing the part with the lowest concentration touch the agar first. Plates were incubated at 37°C inside the anaerobic chamber, at least overnight and longer depending on the species-specific growth requirements. After formation of a symmetrical inhibition ellipse, plates were taken out of the chamber and imaged under controlled lighting conditions (spImager S&P Robotics Inc.) using an 18 megapixel Canon Rebel T3i (Canon Inc. USA). MICs were directly determined from the strip scale at the point where the edge of the inhibition ellipse intersects the MIC test strip. All MICs were determined in duplicates. In cases of an eight-fold difference between the two values, a third replicate was done. In all cases, this resulted in a clear outlier (> 8-fold different from other two MICs) that was removed from the dataset.

### MIC comparison to ChEMBL and EUCAST databases

Previously known MICs were extracted from the ChEMBL database (version 24)^27^ and EUCAST (obtained on May 14, 2018)^24^. Antibiotics from these two datasets were mapped to our dataset by name. Species were mapped using NCBI Taxonomy Identifiers and species names. For MICs from ChEMBL, a keyword-based approach was used to exclude experiments on species with mutations, deletions, insertions, etc. The EUCAST database contains a large number of reported MICs for each compound–species pair. We collapsed these to a single value by calculating the median MIC.

Estimates on the abundance and prevalence of species in the healthy human gut microbiome were calculated using mOTUs v2^48^ as follows: Relative species abundances were determined in 727 shotgun metagenomic samples from donors in the control groups of multiple studies from various countries and continents^49-53^. Prior to taxonomic profiling, metagenomes were quality controlled using the MOCAT2 -rtf procedure^54^, which removed reads with ≥95% sequence identity and an alignment length of ≥45bp to the human genome hg19. Taxonomic profiles were then created using mOTUs version 2.1.0^48^ with parameters -l 75; -g 2; and -c. Afterwards relative abundances below 10^−4^ were set to zero and species with nonzero abundance in <5 samples discarded. For the retained 1,350 species, prevalence was defined as the percentage of samples with nonzero abundance; a prevalence cut-off of 1% was chosen to classify species into “rare” and “common” species. For all species in the MIC dataset, we manually assessed their status as pathogenic or non-pathogenic species using encyclopaedic and literature knowledge. Pathogenic species that occur in more than 1% of healthy people (i.e. are designated as “common”) were classified as “potentially pathogenic species” that can, for example, cause diseases in immunocompromised patients.

### Killing curves and survival assay

Cells were precultured as described in the *growth conditions* section before being diluted to an OD_578_=0.01 and grown for 2 h at 37°C (unless specified otherwise). Next, cells were diluted 1/2 in MGAM containing a 10-fold MIC of erythromycin, azithromycin or doxycycline (final antibiotic concentration is 5-fold MIC) and incubated in the presence of the antibiotic for 5 h at 37°C. At several time-points (0, 1h, 2h, 3h, 4h, 5h), 100 μl of cells were serial-diluted in PBS (10^−1^ to 10^−8^ dilutions) and plated on MGAM-Agar plates for CFU counting. When no cells were detected using this method, a bigger volume of culture (up to 2 ml) was plated to be able to detect CFUs. Agar plates were incubated overnight at 37°C and colonies were counted the next day, either manually, for low CFU numbers, or using the *Analyze Particles* tool from ImageJ^55^.

### Live/dead staining

Cells were precultured as described in the *growth conditions* section before being diluted to an OD_578_=0.01 and grown for 2 h at 37°C. Cells were next diluted 1/2 in MGAM containing 10-fold MIC of erythromycin, azithromycin or doxycycline (final concentration is 5-fold the MIC) and incubated in the presence of the antibiotic for 5 h at 37°C. Then, cells were live/dead stained using the *LIVE/DEAD BacLight Bacterial viability and counting kit* (#L34856 Molecular Probes, ThermoFisher) according to the manufacturer’s protocol before and after antibiotic treatment.

### Flow cytometry

Stained cells were counted using a BD LSRFortessa^TM^ flow cytometer. The forward and side scatter signals (488 nm) as well as the green and red fluorescent signals (488-530/30A filter and 561-610/20A filter, respectively) were acquired. The FSC/SSC detectors were set to logarithmic scale. The flow rate varied between 12 µl/min and 60 µl/min depending on the concentration of each sample, and the analysis was stopped when 10,000 target events were measured. Graphs were generated using the FlowJo V10.3 software (Treestar).

### Microscopy

For live/dead imaging, stained cells were washed twice in 0.85% NaCl before being spotted on 0.85% NaCl +1% agarose pads between a glass slide and a coverslip. For time-lapse imaging, cells were precultured as described in the *growth conditions* section. Cells were then diluted to an OD_578_=0.01 and grown for 3 h at 37°C before being spotted on MGAM +1% agarose pads, supplemented or not with 15 µg/ml erythromycin (5-fold MIC) between a glass slide and a coverslip. Slides were sealed with valap (to avoid/delay oxygen permeation) and taken outside of the anaerobic chamber for imaging. In these conditions, untreated bacteria kept growing rapidly (Movie 1 + 3). The imaging was performed using a Nikon Eclipse Ti inverted microscope, equipped with a Nikon DS-Qi2 camera, a Nikon Plan Apo Lambda 60X oil Ph3 DM phase contrast objective and a Nikon HC mCherry filter set (Ex 562/40; DM 593; BA 641/75) to detect propidium iodide fluorescence. Images were acquired with the NIS-Elements AR4.50.00 software and processed with Fiji v.2.0.0-rc-68/1.52h^56^.

### Growth curves

Cells were precultured as described in the *growth conditions* section. Then, cells were diluted to an OD_578_=0.01 in a 96-well plate sealed with a breathable membrane (Breathe-Easy®) and grown for 2 h. Next, erythromycin was added to the culture to a final concentration of 15 µg/ml (5-fold MIC) and growth curves were acquired for 20 h using a microplate spectrophotometer (EON, Biotek) by measuring the OD_578_ every hour after 30 sec of linear shaking.

### Screen for microbiome-protective antibiotic antagonism

#### Preparation of screening plates

The Prestwick Chemical Library was purchased from Prestwick Chemical Inc. and drugs were re-arrayed, diluted and stored in 96 well format as described before^4^. We prepared drug plates (2 x drug concentration) in MGAM medium and stored them at -30°C. For each experiment, drug plates were thawed, supplemented with the respective antibiotic solution (freshly prepared in MGAM) and pre-reduced in the anaerobic chamber overnight. All rearranging and aliquoting steps were done using the Biomek FXP (Beckman Coulter) system.

#### Inoculation and screening conditions

Strains were grown twice overnight, the second overnight culture was diluted in MGAM to reach OD_578 nm_ 0.04 (4 x the desired starting OD). 25 µl of the diluted cultures were used to inoculate wells containing 50 µl of 2x concentrated Prestwick drug and 25 µl of the 4x concentrated antibiotic using the semi-automated, 96-well multi-channel pipette ep*Motion*96 (Eppendorf). Each well contained 1% DMSO, 20 µM of the Prestwick drug and a species-specific antibiotic concentration that was just inhibitory for the respective strain (0.625 µM for erythromycin, 0.04 µM doxycycline for *B. uniformis* and 0.08 µM doxycycline for *B. vulgatus*). Plates were sealed with breathable membranes (Breathe-Easy®) and OD_578_ was measured hourly after 30 sec of linear shaking with a microplate spectrophotometer (EON, Biotek) and an automated microplate stacker (Biostack 4, Biotek) fitted inside a custom-made incubator (EMBL Mechanical Workshop). Growth curves were collected up to 24 h. For each antibiotic, each species was screen in biological duplicates. All experiments included control wells of unperturbed growth (32 wells per run) and control wells for growth in the presence of the antibiotic only (8 wells per plate).

#### Analysis pipeline and hit calling

All growth curves within a plate were truncated at the transition time from exponential to stationary phase and converted to normalized AUCs using in-run control wells (no drug) as described before^4^. We then calculated z-scores based on these normalized AUCs, removed replicates with 8-fold differences in z-scores to eliminate noise effects, computed mean z-scores across the two replicates and selected combinations with mean z-scores > 3. This selection included 19 potential antibiotic antagonists and we followed up on 14 of them (7 potential erythromycin and 7 potential doxycycline antagonists in either *B. vulgatus* or *B. uniformis* – see Extended Data Fig. 9) in independent experiments. *Validation of microbiome-protective antagonists*. First, we kept the erythromycin/doxycycline concentration constant (0.625 µM for erythromycin, 0.078 µM (*B. vulgatus*)/ 0.039 µM (*B. uniformis*) for doxycycline) and tested concentration gradients of the potential antagonists with ranges depending on the antagonist’s solubility. Compounds were purchased from independent vendors (Suppl. Table 5) and dissolved at 100x starting concentration in DMSO. Eight 2-fold serial dilutions were prepared in 96-well plates with each row containing a different antagonist, sufficient control DMSO wells and wells with just the respective antibiotic (‘antibiotic-only’ control). These master plates were diluted in MGAM medium (50 µl) to 2 x assay concentration and 25 µl freshly prepared antibiotic solution (4x test concentration) was added. Plates were pre-reduced overnight in an anaerobic chamber and inoculated with 25 µl of overnight cultures (prepared as described under *Growth conditions*) to reach a starting OD_578_ of 0.01 and 1% DMSO concentration. Growth was monitored hourly for 24 h after 30 sec of linear shaking (as described for the screen^4^). Experiments were performed in biological triplicates. For analysis, growth curves were converted into normalized AUCs (see above). We accounted for residual growth in the presence of the antibiotic by subtracting the median normalized AUCs of the ‘antibiotic-only’ control per plate. We computed medians across triplicates and considered a normalized AUC > 0.25 as concentration-dependent growth rescue by the antagonist.

#### Checkerboard assays for anaerobic commensals

Validated antagonists were further investigated in 8×8 checkerboard assays, where both antibiotics and antagonists were titrated against each other. Such assays were first performed for the commensals that were originally screened (i. e. *B. vulgatus* and *B. uniformis* – 4 replicates) and later expanded towards six further gut microbes (*B. caccae, B. fragilis* NT, *B. ovatus, B. thetaiotaomicron, P. copri, P. distasonis* – 2 replicates). For vertical gradients, 2-fold serial dilutions of the antagonists were prepared first in 100x in DMSO and diluted in MGAM as described above (section ‘Validation of microbiome-protective antagonists’). Horizontal antibiotic dilution series were freshly prepared in MGAM at 4x final concentration in equidistant concentration steps. Both, vertical and horizontal dilution series were combined (50 µl of the antagonist gradients (2x) and 25 µl of the antibiotic gradients (4x)) in 96 well plates. Plates were pre-reduced under anaerobic conditions overnight, inoculated with 25 µl of diluted overnight culture (at 4x starting OD) and sealed with breathable membrane (Breathe-Easy®). Bacterial growth was monitored once per hour for 24 h after 30 sec linear shaking (Eon + Biostack 4, Biotek) under anaerobic conditions. Growth curves were converted into normalized AUCs as described using in-plate controls to define unperturbed growth.

#### Checkerboard assays for pathogens under aerobic conditions

For three pathogens (*S. aureus* DSM20231 ATCC 12600 and *E. faecium* ATCC19434) 8×8 checkerboard assays were performed in transparent 384 well plates (Greiner BioOne GmbH), with each well containing a total volume of 30 µl in total for *S. aureus* and 55 µl for *E. faecium. S. aureus* strains were grown in tryptic soy broth (TSB, Sigma Aldrich), *E. faecium* in BHI medium (Sigma Aldrich). Drugs were arrayed in 2-fold serial dilutions for the checkerboards. Cell were inoculated at initial OD_595nm_ ∼0.01 from an overnight culture. Plates were sealed with breathable membranes (Breathe-Easy), incubated at 37°C (Cytomat 2, Thermo Scientific) with continuous shaking and OD_595nm_ was measured every 30 min for 16 h in a Filtermax F5 multimode plate reader (Molecular Devices). For *S. pneumoniae* D39, we only tested concentration gradients of the potential antagonists in a constant antibiotic concentration (0.2 µM erythromycin) in BHI medium. All experiments were done at least in 2 biological replicates and 2 technical replicates. Wells in which there was significant condensation were removed and background due to medium was subtracted. Growth curves were trimmed at the transition to stationary phase (9 h for *S. aureus*, 12 h for *E. faecium*). AUCs were calculated and normalised by the median of the internal no-drug control wells (n = 6). Interactions were quantified according to the Bliss interaction model^57^. Interactions were called antagonistic if the median of all the interaction scores for a given checkerboard was above 0.05, synergistic if the value was below -0.05 and neutral if lying between these two cut-offs.

### Phylogenetic analysis/phylogenetic tree construction

In order to generate a phylogenetic tree for the different isolates, the nucleotide sequences for a set of universally occurring, protein coding, single copy phylogenetic marker genes^48,58^ were extracted from reference genomes or genome assemblies using fetchMG^58^ (https://motu-tool.org/fetchMG.html). Within the framework of the ete3 toolkit^59^, ClustalOmega^60^ was used to create sequence alignments for each marker gene independently and all columns with more than 10% gaps were removed. The individual alignments were concatenated and finally, a phylogenetic tree was inferred from the combined alignment using IQTree^61^.

## Data availability

Data is available upon request.

## Code availability

Code is available upon request.

## SUPPLEMENTARY INFORMATION

### Supplementary tables

**Supplementary Table 1**

Annotation and classification of antibiotics

**Supplementary Table 2**

MICs strips used in this study

**Supplementary Table 3**

MICs as determined by MIC strips for selected antibiotics

**Supplementary Table 4**

Hitlist: Screen for microbiome-protective antagonists to erythromycin/tetracycline

**Supplementary Table 5**

Compounds used in this study

### Supplementary movies

**Movie 1**

Time-lapse of *B. vulgatus* growing on MGAM-Agarose 1% pad.

**Movie 2**

Time-lapse of *B. vulgatus* growing on MGAM-Agarose 1% pad containing 5-fold MIC of erythromycin.

**Movie 3**

Time-lapse of *B. uniformis* growing on MGAM-Agarose 1% pad.

**Movie 4**

Time-lapse of *B. uniformis* growing on MGAM-Agarose 1% pad containing 5-fold MIC of erythromycin.

## Acknowledgements

We thank Stephan Göttig and the Typas lab for feedback on the manuscript. We thank Ana Rita Brochado, Sarela Santamarina and the EMBL Flow Cytometry Core Facility for help with experimental design. We acknowledge EMBL and the JPIAMR grant Combinatorials for funding. LM and MP were supported by the EMBL Interdisciplinary Postdoc (EIPOD) program under Marie Sklodowska Curie Actions COFUND (grant 291772) and LM is now supported by the CMFI Cluster of Excellence (EXC 2124). CG is recipient of an EMBO long-term postdoctoral fellowship and an add-on fellowship from the Christiane Nüsslein-Volhard-Stiftung. AT is supported by ERC consolidator grant uCARE.

## Author contributions

This study was conceived by KRP, PB and AT; designed by LM, CVG, MP and AT; and supervised by LM and AT. Experiments were conducted by LM, CVG and MP. TB and EEA contributed to MIC measurements and EC to checkerboard analyses. Data preprocessing, curation and comparisons to existing databases were performed by JW, MK, AM, UL, SKF and GZ. Data interpretation was performed by LM, CVG, JW, GZ and AT. LM, CVG and AT wrote the manuscript with feedback from all authors; LM, CVG, JW and MK designed figures with inputs from GZ and AT. All authors approved the final version for publication.

## Author information

EMBL has filed a patent application on using the antidotes identified in this study for prevention and/or treatment of dysbiosis and for microbiome protection (European patent application number EP19216548.8). LM, CVG, EC and AT are listed as inventors. Correspondence and requests for materials should be addressed to.

---

**Extended Data Figure 1.**
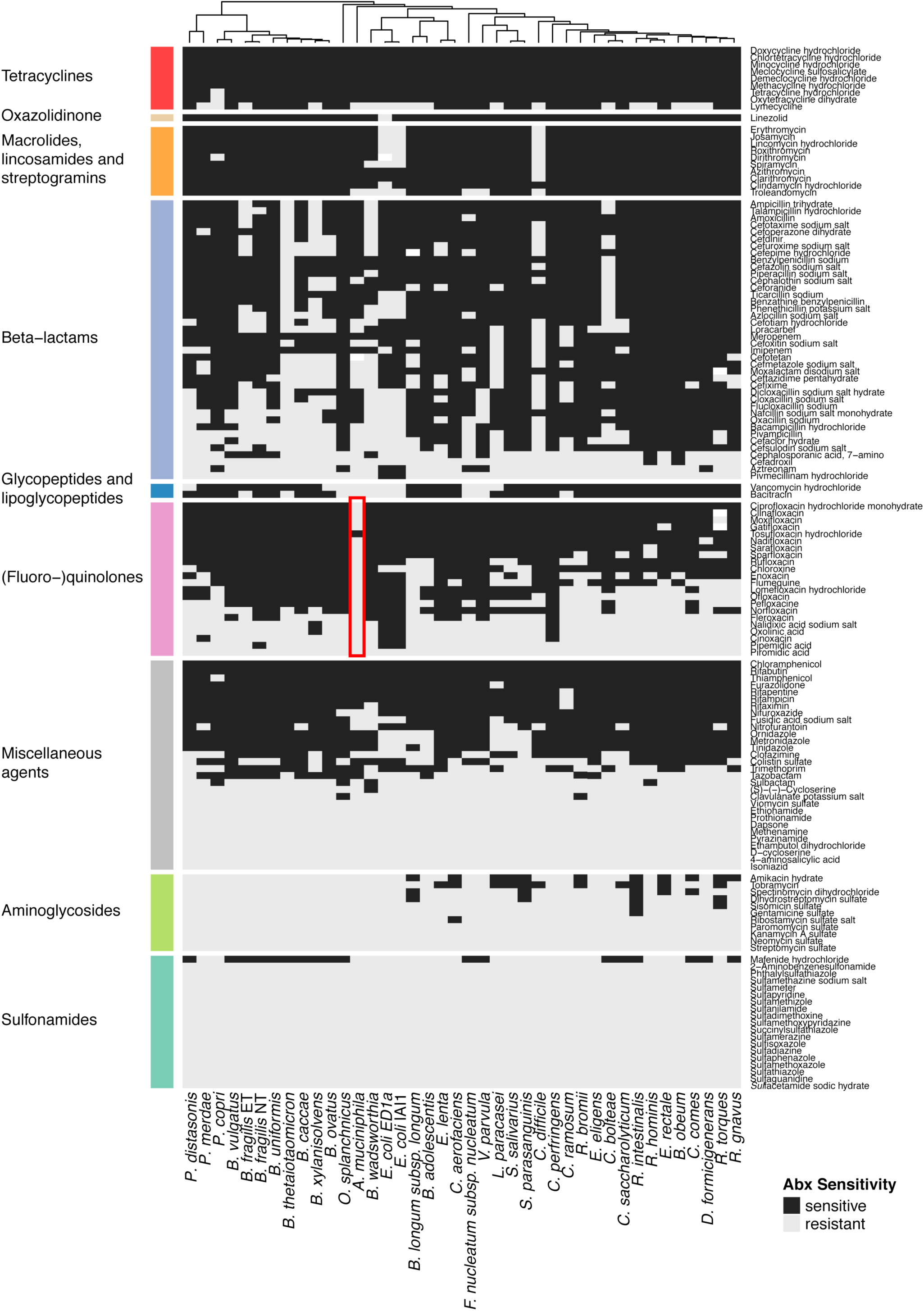
Effects of 144 antibiotics on 40 human gut commensals. Heat map according to sensitivity or resistance of each strain to the respective antibiotic at a concentration of 20 µM. Antibiotics are grouped according to drug classes and species are clustered according to their responses across the 144 antibiotics tested. Data is replotted from^4^. *Akkermansia muciniphila* (Muc, DSM22959, type strain) is resistant to nearly all quinolone antibiotics (red box). We consolidated this finding by MIC determination for Ciprofloxacin (>32 µg/ml), Gatifloxacin (>32 µg/ml), Moxifloxacin (>32 µg/ml), Norfloxacin (>256 µg/ml) and Ofloxacin (>32 µg/ml).

**Extended Data Figure 2.**
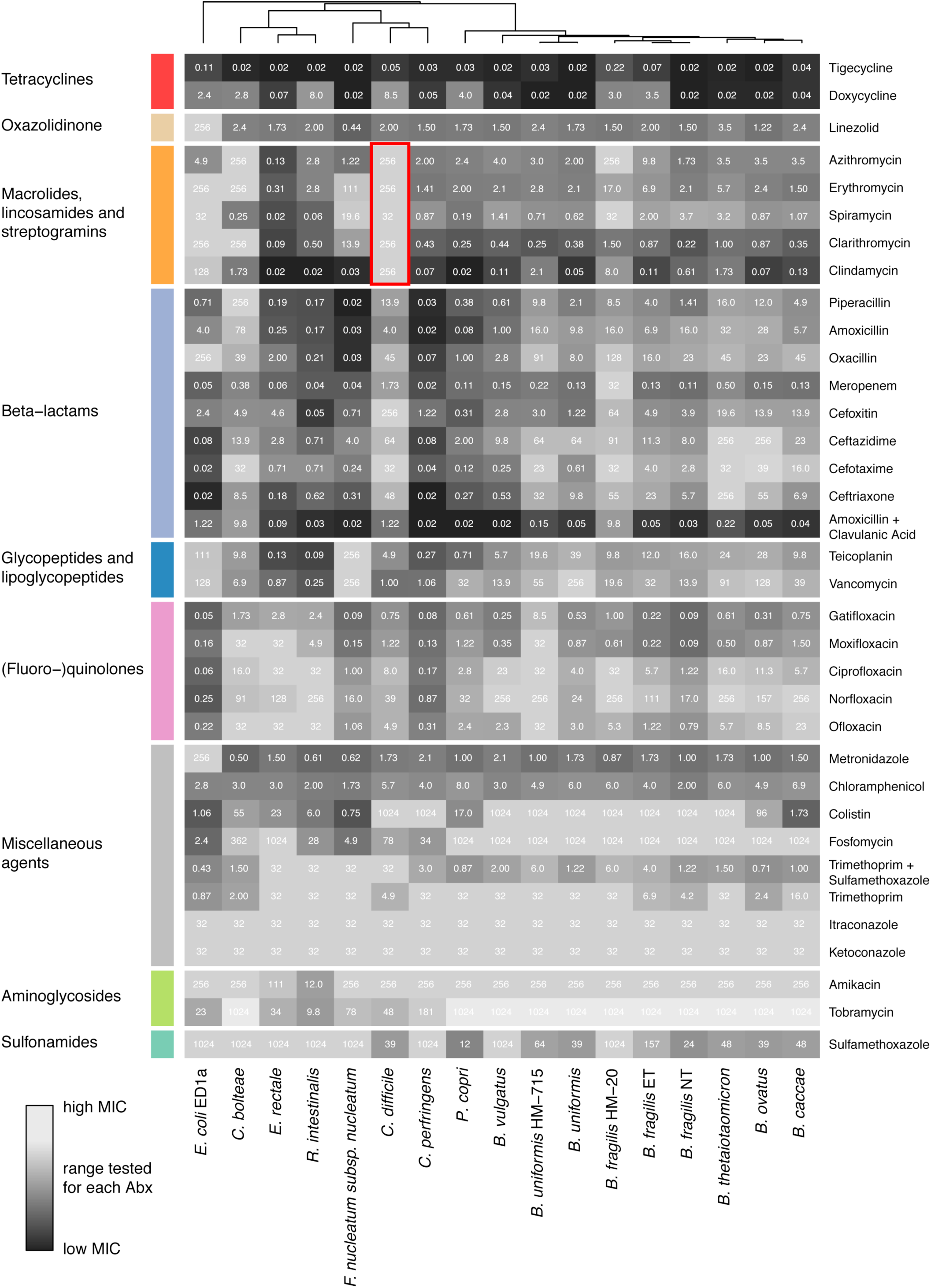
MICs for 17 species on 35 antimicrobials. Heat map depicts MICs for each drug-strain pair in µg/ml. Heat map color gradient is adjusted to the MICs concentration range tested on the respective MIC test strip. Black depicts sensitivity and light grey indicates resistance. Mean values across two biological replicates are shown (Suppl. Table 3). *C. difficile* is particularly resistant to all tested macrolides and clindamycin (red box).

**Extended Data Figure 3.**
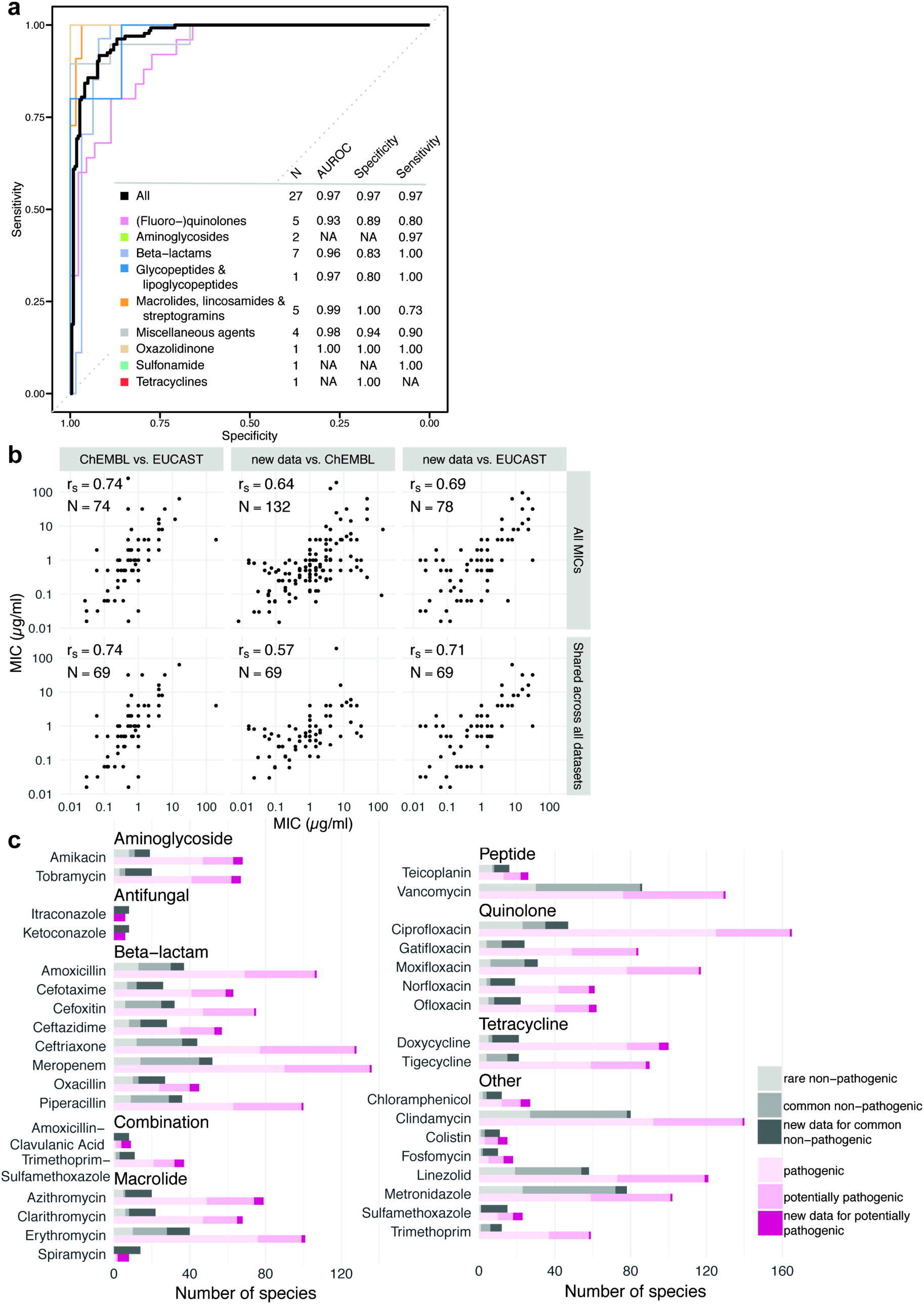
MIC dataset validates antibiotic sensitivity profiles from the screen dataset and is consistent with publically available MICs. **a**. Receiver operating characteristic (ROC) curve analysis was performed to evaluate sensitivity and specificity of the screen^4^ using the MIC dataset. Results from the screen were considered as validated if MICs were below/above the 20 µM antibiotic concentration that was tested in the screen (allowing a two-fold error margin). N is the number of antibiotics that we tested both in the screen and determined MICs for, AUROC is the area under the characteristic ROC. TN denotes true negatives, FP false positives, TP true positives, FN false negatives. **b**. Comparison including Spearman correlation coefficients of the MICs from this study to MICs from the ChEMBL ^27^ and EUCAST ^24^ databases. Panels in the upper row: comparison between all MICs that are shared between the two indicated datasets. Panels in the lower row: comparison of the 69 MICs that are shared across all three datasets. Despite experimental differences, our MICs correlate well with available EUCAST/ ChEMBL data. **c**. Number of the sum of new (this study) and already available MICs (EUCAST/ ChEMBL) per drug according to antibiotic class and prevalence/virulence of the bacterial species. The new dataset expands MICs across the board and specifically fills the knowledge gap on non-pathogenic species.

**Extended Data Figure 4.**
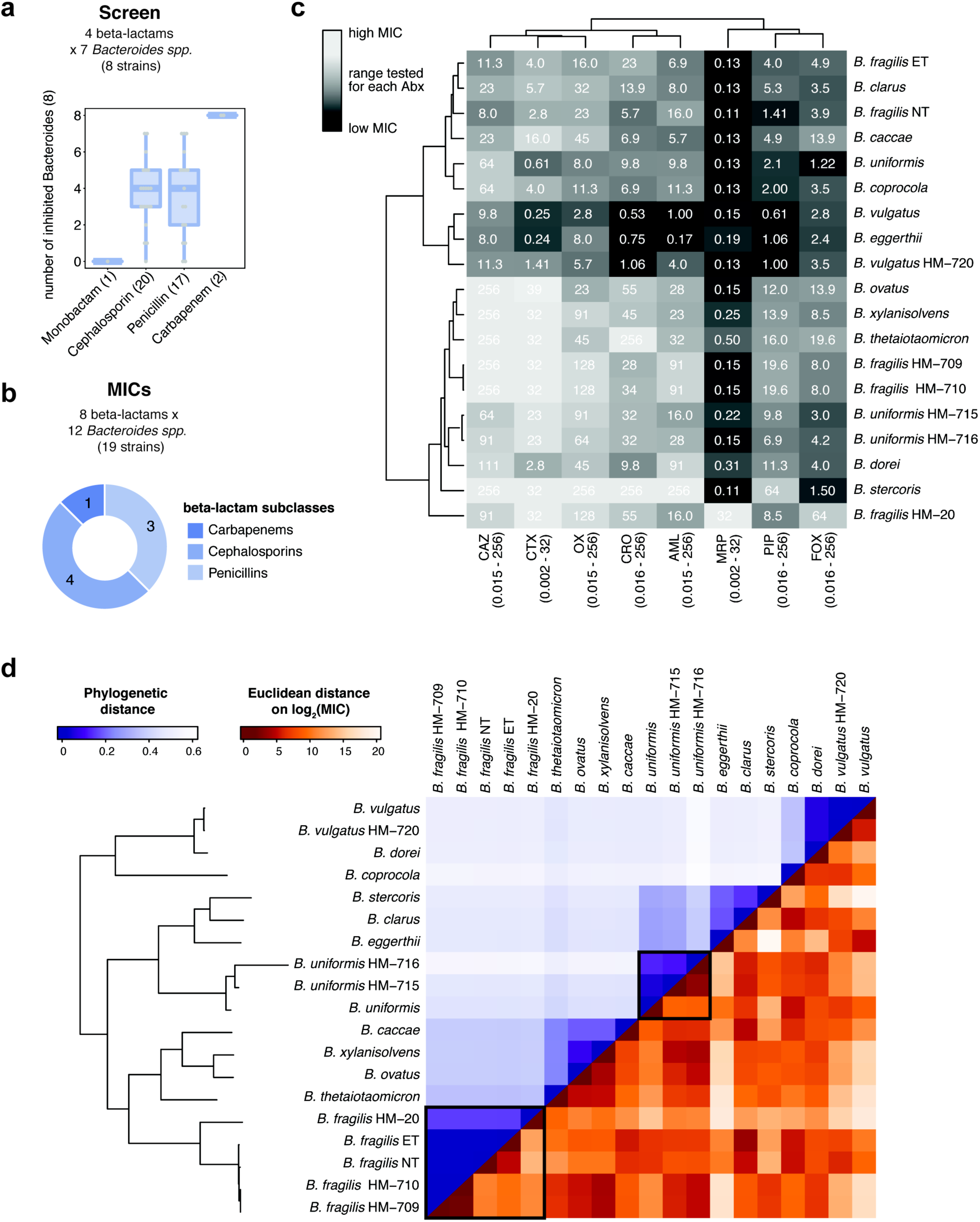
β-lactam antibiotic resistance profiles do not recapitulate phylogenetic relationship between *Bacteroides spp*. **a**. Number of inhibited *Bacteroides spp*. (out of 8 tested) at 20 µM per β-lactam subclass, based on the initial screen^4^. Number of drugs per class tested are shown in parenthesis. Boxes plotted as in Figure 1d. **b**. Overview of the number of drugs tested per β-lactam subclasses on *Bacteroides spp*.; compared to ED Figure 2, 10 additional strains were tested: *B. eggerthii, B. clarus, B. coprocola, B. vulgatus* HM-720, *B. xylanisolvens, B. fragilis* HM-709, *B. fragilis* HM-710, *B. uniformis* HM-716, *B. dorei* and *B. stercoris*. **c**. MIC heat map for 8 β-lactam antibiotics on 19 *Bacteroides spp*. Strains are clustered according to resistance profiles across all β-lactam antibiotics, drugs are clustered according their effects on *Bacteroides spp*. MICs values are based on two biological replicates and are partially replotted from Extended Data Fig. 2. Heat map gradients are adjusted to the antibiotic concentration ranges tested with lighter color depicting resistance and darker color depicting sensitivity. **d**. Heat map of phylogenetic relationship between *Bacteroides spp* (upper triangular matrix) ordered by phylogeny and their resistance profiles across β-lactam antibiotics (lower triangular matrix). Colors represent the pairwise phylogenetic distance and the Euclidean distance on the log2 transformed MICs for β-lactams (panel c). Examples of strains from the same species (*B. fragilis* / *B. uniformis*) that respond differently to β-lactam antibiotics, are highlighted.

**Extended Data Figure 5.**
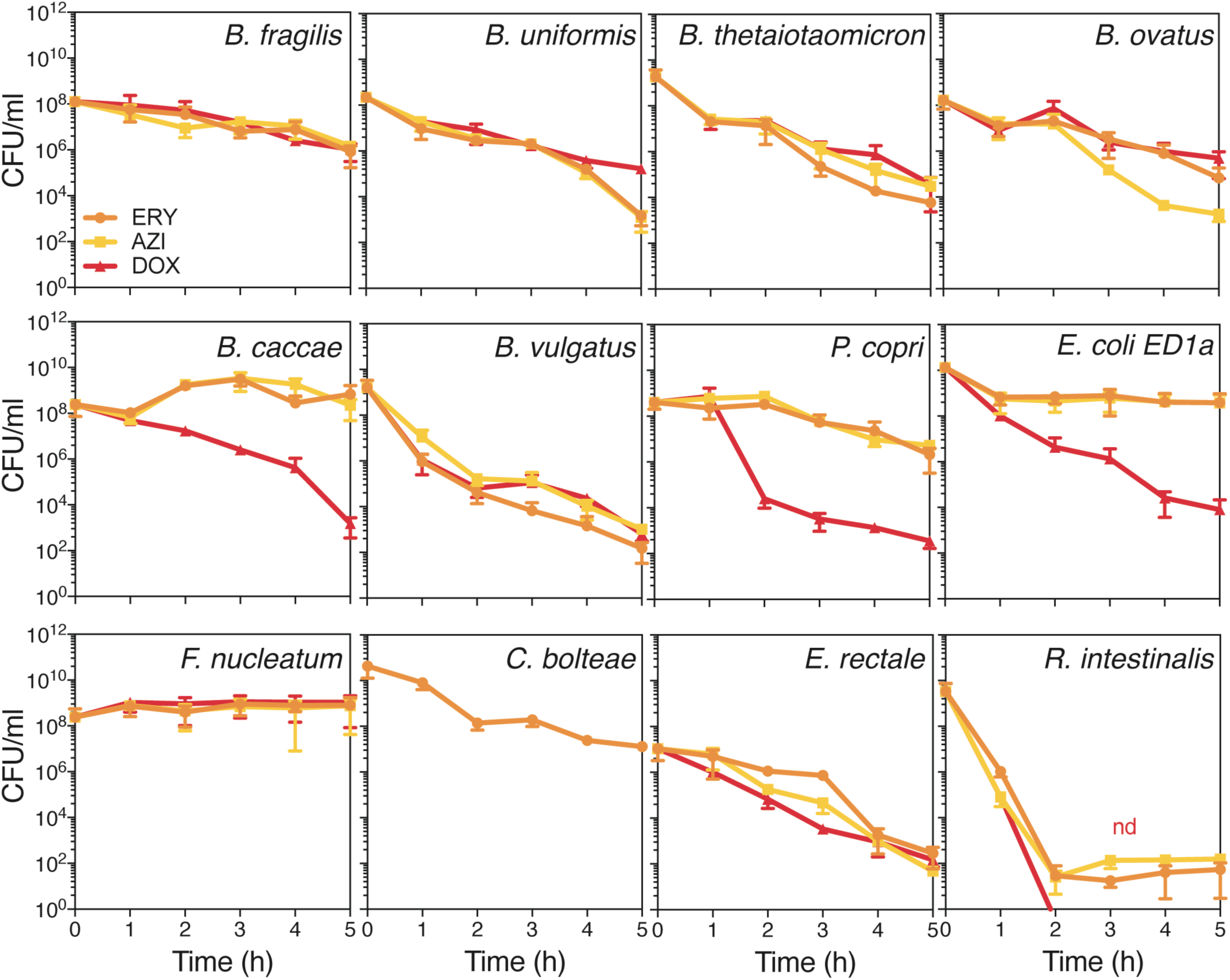
Time-kill curves of 12 abundant gut microbes after treatment with macrolides and tetracyclines. Survival of 12 abundant gut microbes was assessed by CFU counting over a 5 hour-treatment of either ERY, AZI or DOX. This graph shows the mean±SD of 3 independent experiments.

**Extended Data Figure 6.**
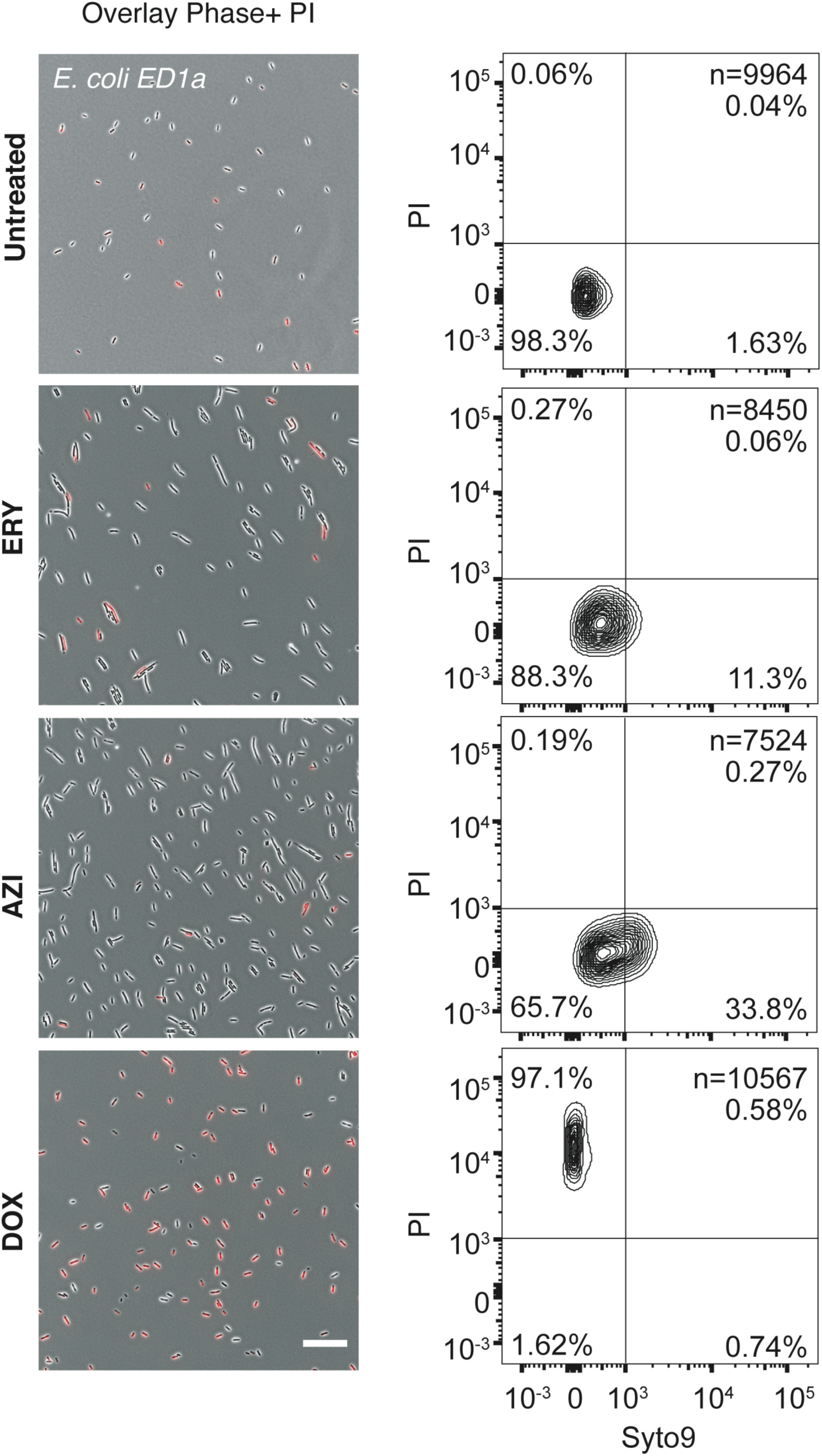
Live/dead staining of macrolide or tetracycline-treated *E. coli* ED1a. The left panel shows an overlay of phase contrast and fluorescence microscopy images of propidium iodide (PI)-stained *E. coli* ED1a before and 5 hours after ERY, AZI or DOX treatments. The number of cells on each frame has no meaning, as cultures were concentrated before imaging; the scale bar is 10 µM. The right panel shows the corresponding quantification of live/dead-stained cells by flow cytometry with Syto9 on the x-axis (live cells) and PI on the y-axis (dead cells). Both the total number of measured events (n) and the percentage of cells found in each quadrant are indicated on the graphs.

**Extended Data Figure 7.**
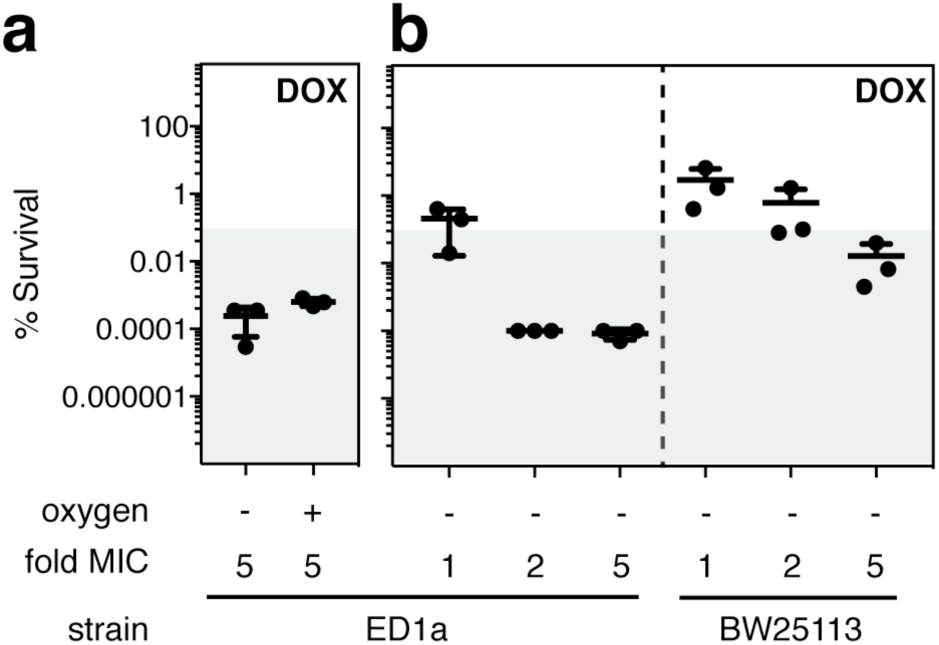
Effect of oxygen and strain specificity on survival after doxycycline treatment. **a**. The survival of *E. coli* ED1a was assessed after a 5-hour treatment with 5-fold MIC of DOX in the presence or absence of oxygen. Killing was similarly effective in both conditions. **b**. The survival of *E. coli* ED1a and *E. coli* BW25113 were assessed after a 5-hour treatment with 1, 2 and 5-fold MIC of DOX in MGAM medium in anaerobic conditions. The lab strain is more resistant to killing with doxycycline becoming boarder-line bactericidal at higher MICs.

**Extended Data Figure 8.**
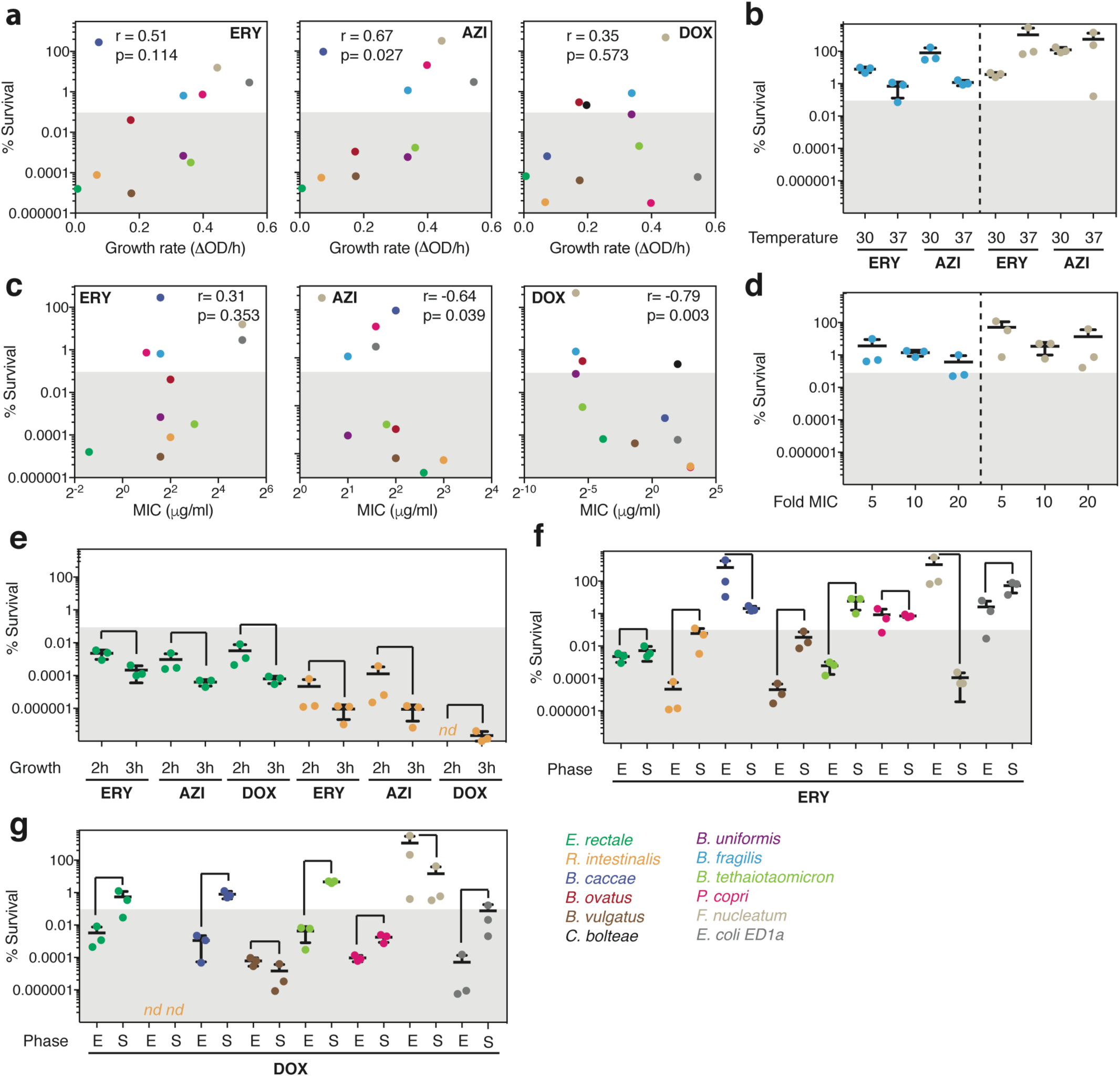
Assessing potential confounding factors for the killing capacities of erythromycin, azithromycin and doxycycline. **a**. Scatter plot of individual bacterial growth rates and percentage survival after a 5-hour treatment with 5-fold MIC of ERY, AZI or DOX treatments. *r* indicates the Spearman correlation coefficient. Tested species are color-coded here and in all panel thereafter as indicated in the bottom of this figure. Positive correlations for macrolides were tested further in **b** to check if changing growth rate in same species affects percentage killed. **b**. The survival of *B. fragilis* (blue) and *F. nucleatum* (beige) were assessed after a 5-hour macrolide treatment (5-fold MIC of ERY and AZI) at either 30°C (slow growth) or 37°C (fast growth) to test the effect of slowing down growth on survival. No significant change observed. This graph shows the mean±SD of three independent experiments. **c**. Scatter plot of MICs and percentage survival after a 5-hour treatment with 5-fold MIC of ERY, AZI or DOX treatments. *r* indicates the Spearman correlation coefficient. Doxycycline exhibited a strong and significant anti-correlation, that is that species which were more sensitive to doxycycline (lower MIC) were not killed when they were treated with 5-fold MIC concentrations. Thus, we tested further whether increasing the drug concentration in some of those sensitive strains decreased the % of survival (panel **d**). **d**. The survival of *B. fragilis* (blue) and *F. nucleatum* (beige) were assessed after a 5-hour treatment with increasing concentrations of DOX (5, 10 or 20-fold of MIC) to test whether higher concentrations of DOX induced more killing. This seemed not be the case. This graph shows the mean±SD of three independent experiments. **e**. To evaluate whether outgrowth of stationary phase and homogeneity of population affected our results, we selected two slow-growing strains, *E. rectale* and *R. intestinalis* and grew for 2 or 3 hours after being diluted from an overnight culture to an of OD_578_ 0.01. Both strains were then treated for 5 hours with 5-fold MIC of ERY, AZI or DOX and their survival was assessed to test the impact of the growth phase on the percentage survival. Although slight differences were observed and 3h grown cultures were killed more effectively (presumably because more cells had exited stationary phase and were growing exponentially by then), the general trends remained the same. If anything, this means that we are underestimating the killing for slow-growers, since we performed all other experiments with 2 hours outgrowth. This graph shows the mean±SD of three independent experiments. **f**. The survival of 8 selected gut microbes was measured after treating cells in exponential phase (E – 2 hours after dilution from an overnight culture) or in stationary phase (S – overnight growth) with 5-fold MIC of ERY for 5 hours to test the impact of the growth phase on the percentage survival. As expected, survival is higher in stationary phase for half of the strains, but in some cases stationary phase cells were as or more sensitive than exponentially growing cells – this is the case for *B. caccae* and *F. nucleatum*. This graph shows the mean±SD of three independent experiments. **g**. Same as in **f** but with DOX. Similar effects observed as in f, with more than half of strains becoming more resistant in stationary phase.

**Extended Data Figure 9.**
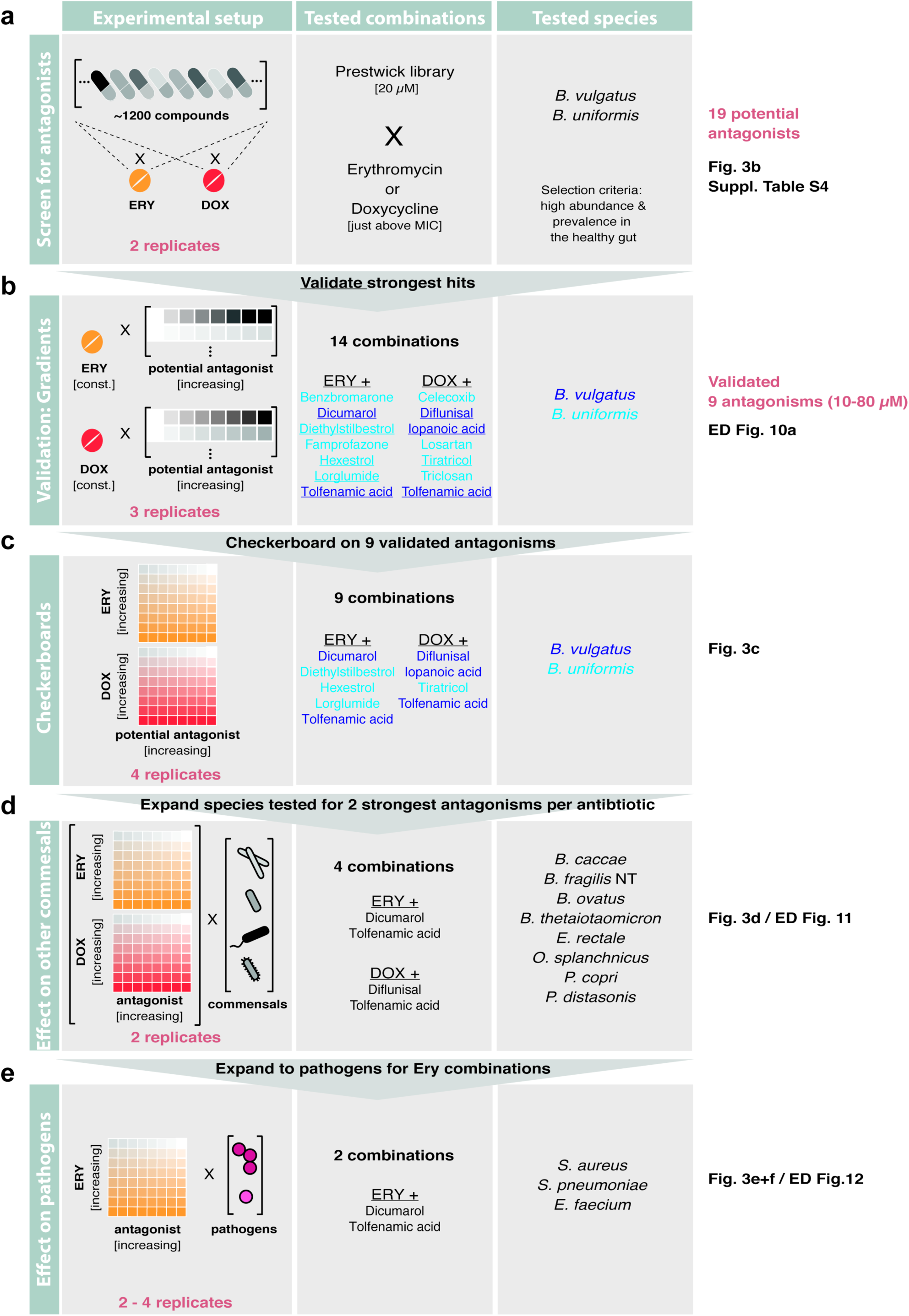
Schematic overview of screen for microbiome-protective antibiotic antagonisms. Workflow with decision process on which antagonist to move on to next evaluation step.

**Extended Data Figure 10.**
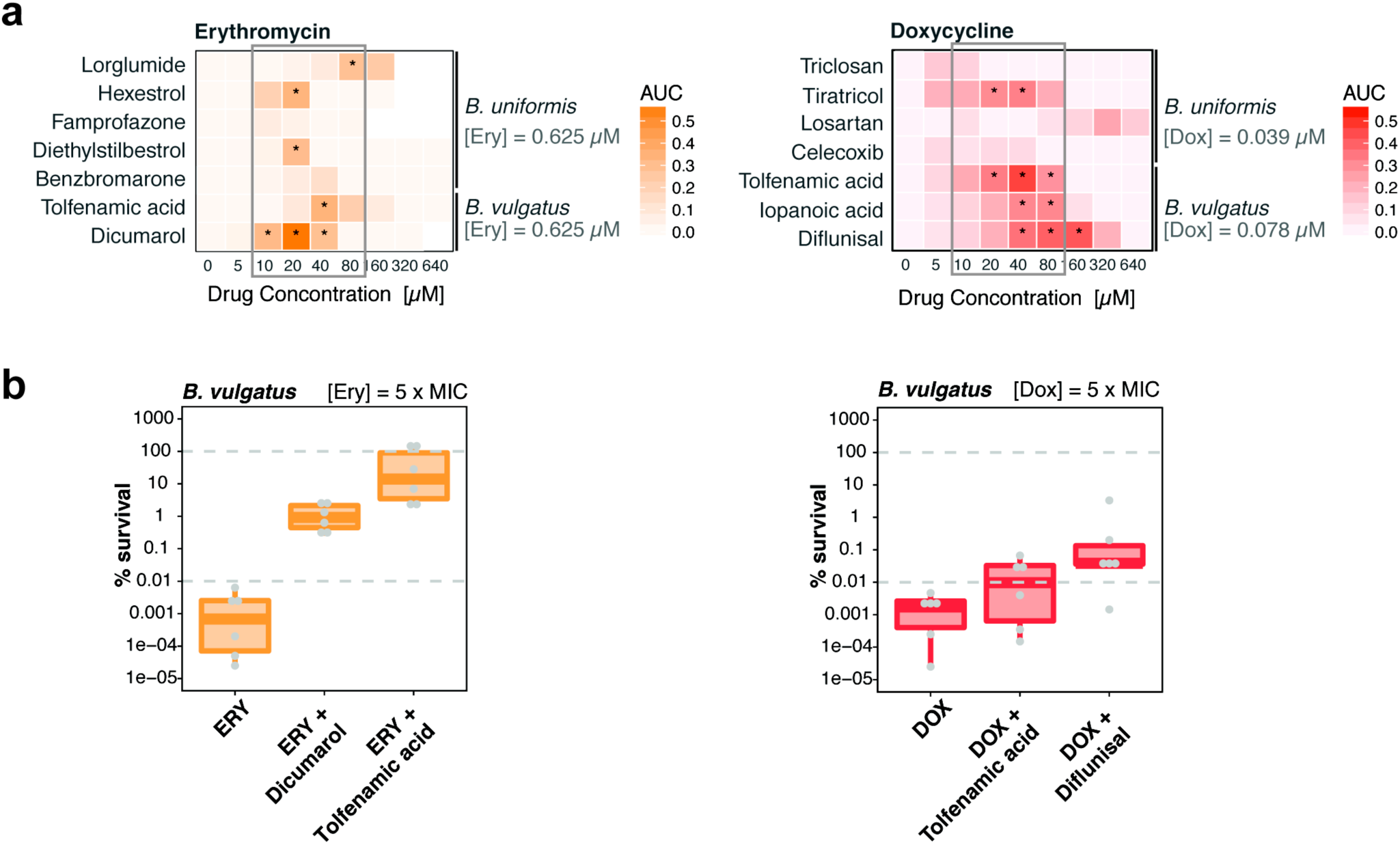
Validation of potential microbiome-protective antagonists. **a**. Validation of the strongest antagonists in independent experiments. Erythromycin and doxycycline concentrations were kept constant ([ERY]=0.625 µM, [DOX] = 0.039 / 0.078 µM) and concentration ranges were tested for antagonist. Asterisks indicate that at least 25% of the bacterial growth (compared to no drug controls) could be rescued by the antagonist at a given concentration. Heat map depicts median AUCs across triplicates. **b**. Percentage of surviving *B. vulgatus* cells were determined after 5 h incubation with either erythromycin (3.25µM) or doxycycline (0.4 µM) alone or in presence of the antagonist dicumarol (20 µM), tolfenamic acid (40 µM) or diflunisal (80 µM). Data is based on 3 independent experiments. Boxplots are plotted as in Figure 1d.

**Extended Data Figure 11.**
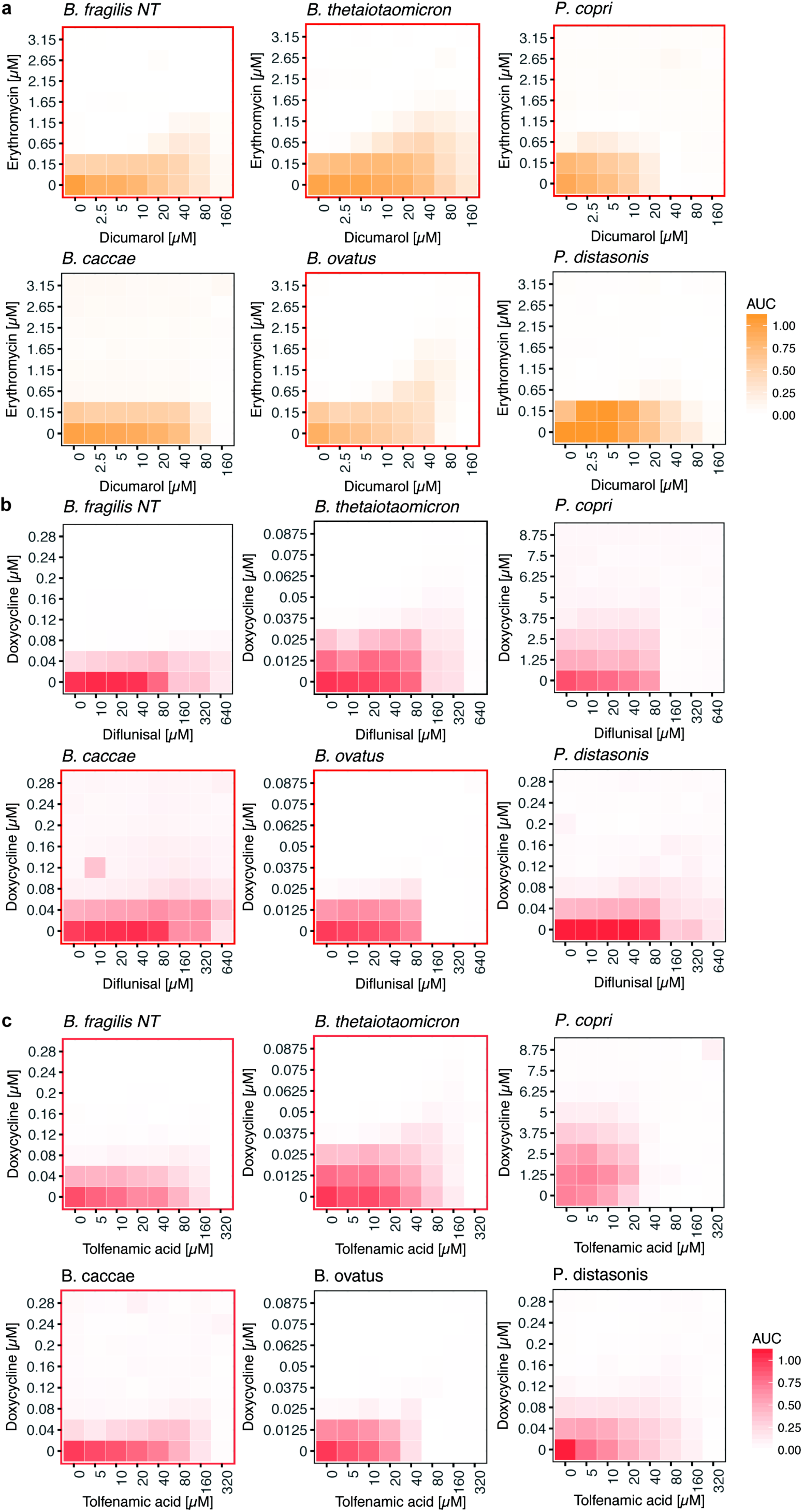
Effect of antidotes on further gut commensals. 8 x 8 checkerboard assays to investigate if antidote is also protective for additional gut commensals for the following combinations: erythromycin and dicumarol (**a**), doxycycline and diflunisal (**b**) and doxycycline and tolfenamic acid (**c**). Heat map depicts bacterial growth based on median AUCs from two independent replicates. Red contours indicate antagonistic drug interactions.

**Extended Data Figure 12.**
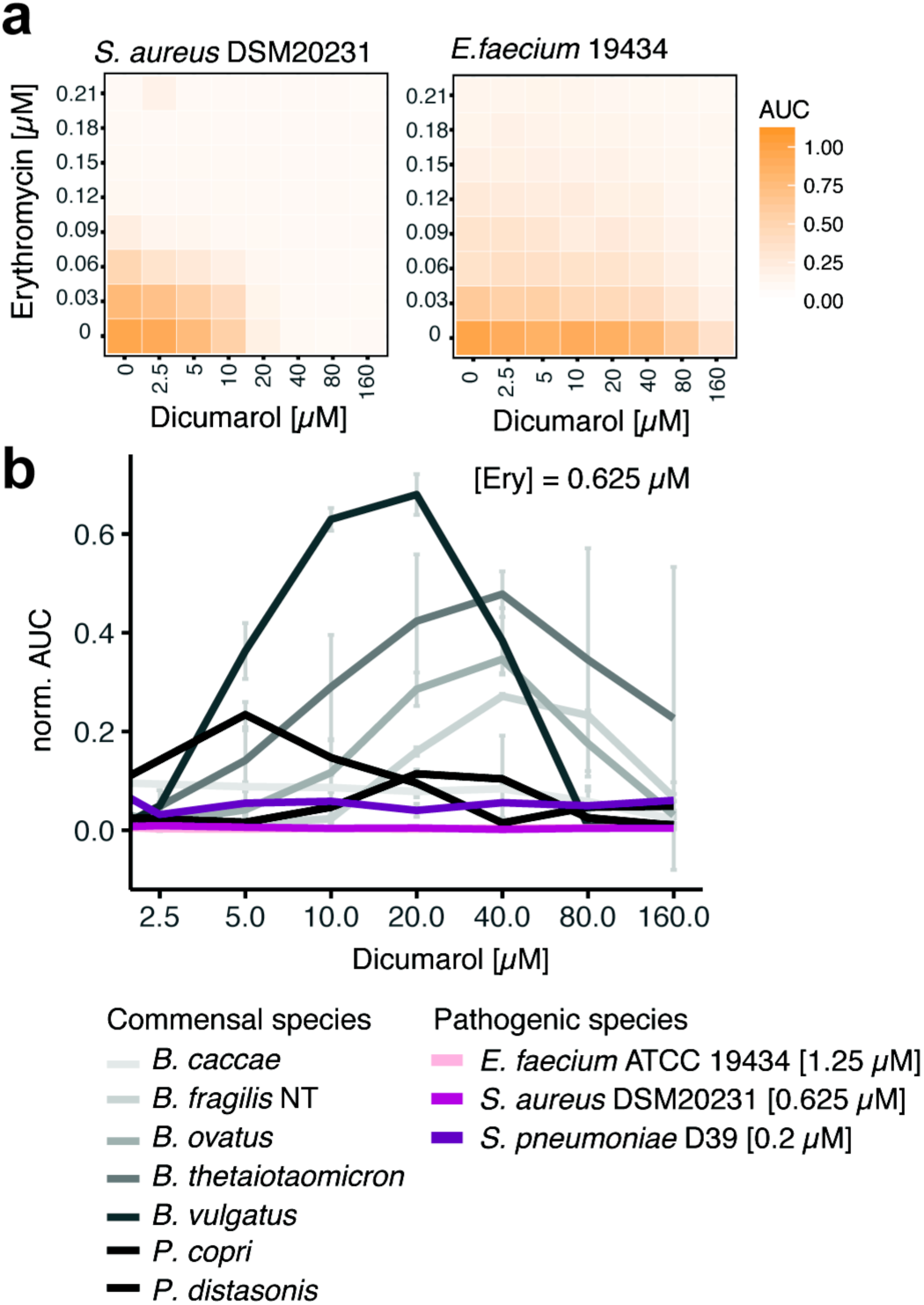
Effect of the antidote dicumarol on pathogens, relatively to commensal species. **a**. Checkerboard assays for the drug combinations erythromycin-tolfenamic acid and erythromycin-dicumarol on the pathogens *S. aureus* (two different strains) and *E. faecium*. Heat map depict median normalized AUCs of checkerboard assays (at least three independent replicates). **b**. Dicumarol rescues commensal growth (based on median AUCs, N=2) at clinical relevant erythromycin concentrations in a concentration-dependent manner. Erythromycin still retains its activity against pertinent pathogens such as *S. aureus, E. faecium* and *S. pneumoniae* and is even slightly more active (synergy) for *E. faecium* (based on median AUCs, N=3). Error bars depict standard deviation.

